# Anisotropic ESCRT-III architecture governs helical membrane tube formation

**DOI:** 10.1101/716308

**Authors:** Joachim Moser von Filseck, Luca Barberi, Nathaniel Talledge, Isabel Johnson, Adam Frost, Martin Lenz, Aurélien Roux

## Abstract

ESCRT-III proteins assemble into ubiquitous membrane-remodeling polymers during many cellular processes. Here we describe the structure of helical membrane tubes that are scaffolded by bundled ESCRT-III filaments. Cryo-ET reveals how the shape of the helical membrane tube arises from the assembly of distinct bundles of protein filaments that bind the membrane with different mean curvatures. Cryo-EM reveals how one of these ESCRT-III filaments engages the membrane tube through a novel interface. Mathematical modeling of the helical membrane tube suggests how its shape emerges from differences in membrane binding energy, positional rigidity, and membrane tension. Altogether, our findings support a model in which increasing the rigidity of ESCRT-III filaments through the assembly of multi-strands triggers buckling of the membrane.

**One Sentence Summary:** ESCRT-III heteropolymers deform membranes into helical tubes.

## Main Text

The Endosomal Sorting Complexes Required for Transport (ESCRT)-III proteins are an evolutionarily ancient family of proteins that execute membrane scission in different cellular contexts (reviewed in (*1*)). ESCRT-III can polymerize into rings and spirals in solution (*2–4*) or on membrane substrates (*5, 6*). When single or several ESCRT-III proteins are incubated with model membranes *in vitro* or over-expressed in cells, they deform membranes into straight and conical tubes (*6, 7*), exemplified in most detail by the formation of tubules by CHMP1B alone and in complex with IST1 (*6*). Similar but inverted conical structures are also observed *in vivo* by overexpression of CHMP4A/B (*7*) and at the neck of budding Gag envelopes (*8*). Mechanistically, we have previously shown that flat spirals formed on lipid membranes from the ESCRT-III protein Snf7 can accumulate elastic energy and that this energy can be channeled to shape a flat membrane into a tube through a buckling transition (*5, 9*). The highly flexible Snf7 polymer alone is unable to deform the membrane because its interaction with the lipid surface is stronger than its tendency to transition from a flat spiral into a helical polymer. Hence, Snf7 spirals fail to deform artificial membranes *in vitro* (*5*). The addition of Vps24/Vps2 to Snf7 flat spirals may trigger the buckling shape transition, however, because they form helical polymers together (*3, 10*). We and others have previously shown that the addition of Vps24/Vps2 leads to the formation a second, parallel strand next to the Snf7 filament (*10, 11*). Thus, buckling may be triggered by the formation of the composite polymer. In this model, assembly of the second strand increases the energy cost of the flat spiral conformation and favor buckling into an energetically preferred helical conformation. However, the shape transition from a flat spiral into a cylindrical helix implies that the filament-membrane or intra-filament interactions are altered (*12*). We set out to determine how yeast Snf7, Vps24, and Vps2 co-assemble multi-stranded polymers capable of deforming lipid bilayers *in vitro*.

To liposomes incubated with recombinant Snf7 and decorated by flat Snf7 spirals (*5*), we added recombinant Vps24 and Vps2 and incubated the mixture for several hours. Using negative stain EM, we observed a mixture of vesicles decorated with flat spirals (Fig. 1A) (*5, 11*) and helical tubes that were decorated with filamentous protein polymers (Fig. 1B, C). Cryo-EM of the flat spirals and the helical tubes confirmed that both structures were organized on the lipid membrane (Fig. 1D-F) and that the regularity of the helical tubes made them amenable to higher-resolution imaging. We thus focused on the helical, tubular membrane protrusions (Fig. 1B, C, E, F) and found that they only form in the presence of all three proteins (Fig. S1A-C). These helical lipid membrane tubes have an average diameter of 23.9 ± 3.7 nm and are coiled into a helix with an outer diameter of 82.3 ± 6.1 nm and a pitch of 53.1 ± 7.6 nm (all values average ± SD) (Fig. S1D-G). Their prevalence increased with incubation time, indicating thermodynamic stability.

**Fig. 1:**
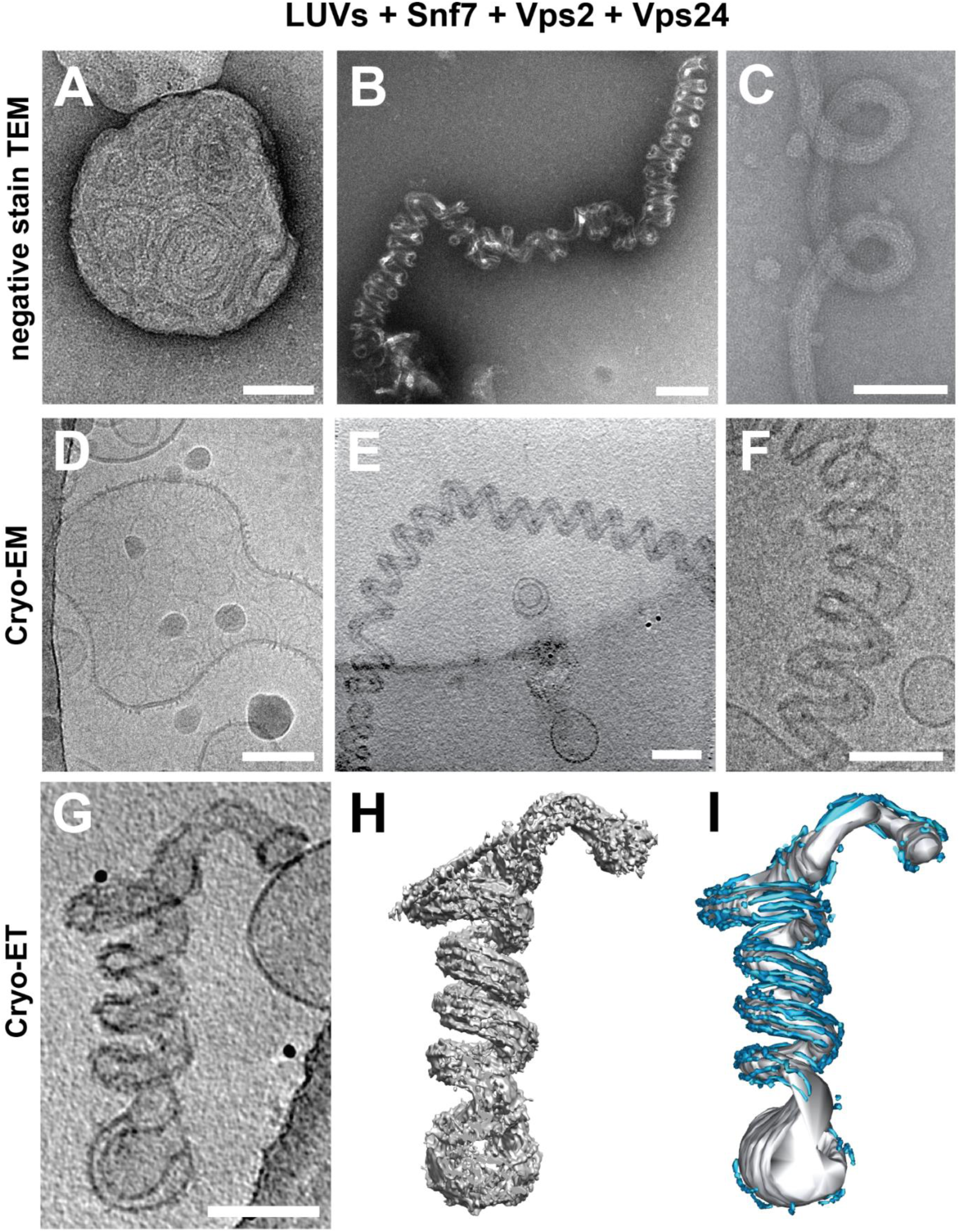
Helical tubulation of liposomes by ESCRT-III heteropolymers. Electron micrographs showing undeformed liposomes (A, D) and helical membrane tubes (B, C, E, F) decorated with Snf7/Vps24/Vps2 on negatively stained (A-C) and vitrified (D-F) samples. The reconstructed cryo-ET volume of a helical membrane tube projected in Z (G) and volume view after filtering (H) or manual segmentation (I) showing the organization of protein filaments (cyan) along the helical membrane tube axis (grey). All scale bars 100 nm.

A helical membrane tube is an unusual shape because it is energetically unfavorable due to the high membrane curvatures. Cylindrical stacks of lipid membranes have been reported to remodel into helical tubes in the presence of specific membrane-binding polymers, although the mechanism of the membrane remodeling was not elucidated (*13*). Helical polymers may stabilize helical membrane tubes by relying on a form of geometric “frustration” that arises when there is an incompatibility between the preferred direction of curvature of the helical polymer and the positioning of the membrane-binding proteins along its length (*14*). We hypothesize that a similar incompatibility determines the preferred helical morphology of tubes in our experiments. Indeed, since Snf7/Vps24/Vps2 form helical filaments, those filaments could wind around a straight tube, binding to it along the direction of their preferred curvature, such as BAR domain-containing protein-coated or dynamin-coated membrane tubes (*15*). Alternatively, they could force the tube to follow their helical path, in this case, being able to bind it in the direction perpendicular to their preferred curvature as well.

To visualize the ESCRT-III filament organization around the helical tubes, we performed cryogenic electron tomography (cryo-ET) on vitrified helical membrane tubes and used image filtering and manual segmentation on reconstructed tomographic volumes. All tubes appeared as left-handed helices, though we cannot confirm this is the handedness without a chiral internal standard. On the surface of the tubes, we observed six to eight filaments parallel to the tube axis forming multi-stranded bundles (Fig. 1G-I, Fig. S1H-J, Movies S1-S2). The filaments are almost always excluded from the inside of the tube helix and have the same thickness as negatively stained, double-stranded Snf7/Vps24/Vps2 heteropolymers (4.9 ± 0.5 nm; average ± SD) (*11*).

To obtain a more detailed view of the filament organization, we performed subtomogram averaging (STA) on slices along the helical tube axis. The variability in tube dimensions in the dataset made it impossible to resolve the entire helical tube. We, therefore, focused on the filaments on the outer tube surface and obtained a reconstruction at 32 Å resolution (Fig. 2). This map revealed that the filaments cluster in three separate regions with two clearly defined grooves between them (Fig. 2A, Fig. S2A). The central cluster, containing two filaments, covered a 13 nm wide region around the equator of the tube (equatorial filaments, blue). Two additional filament clusters, each containing 2-3 filaments, are shifted up and down from the equator, respectively, (polar filaments, red) and appeared wider (16-20 nm) (Fig. 2B-D). The resolution of the shifted, polar filaments was limited as their positions varied more with tube diameter compared to the equatorial region.

**Fig. 2:**
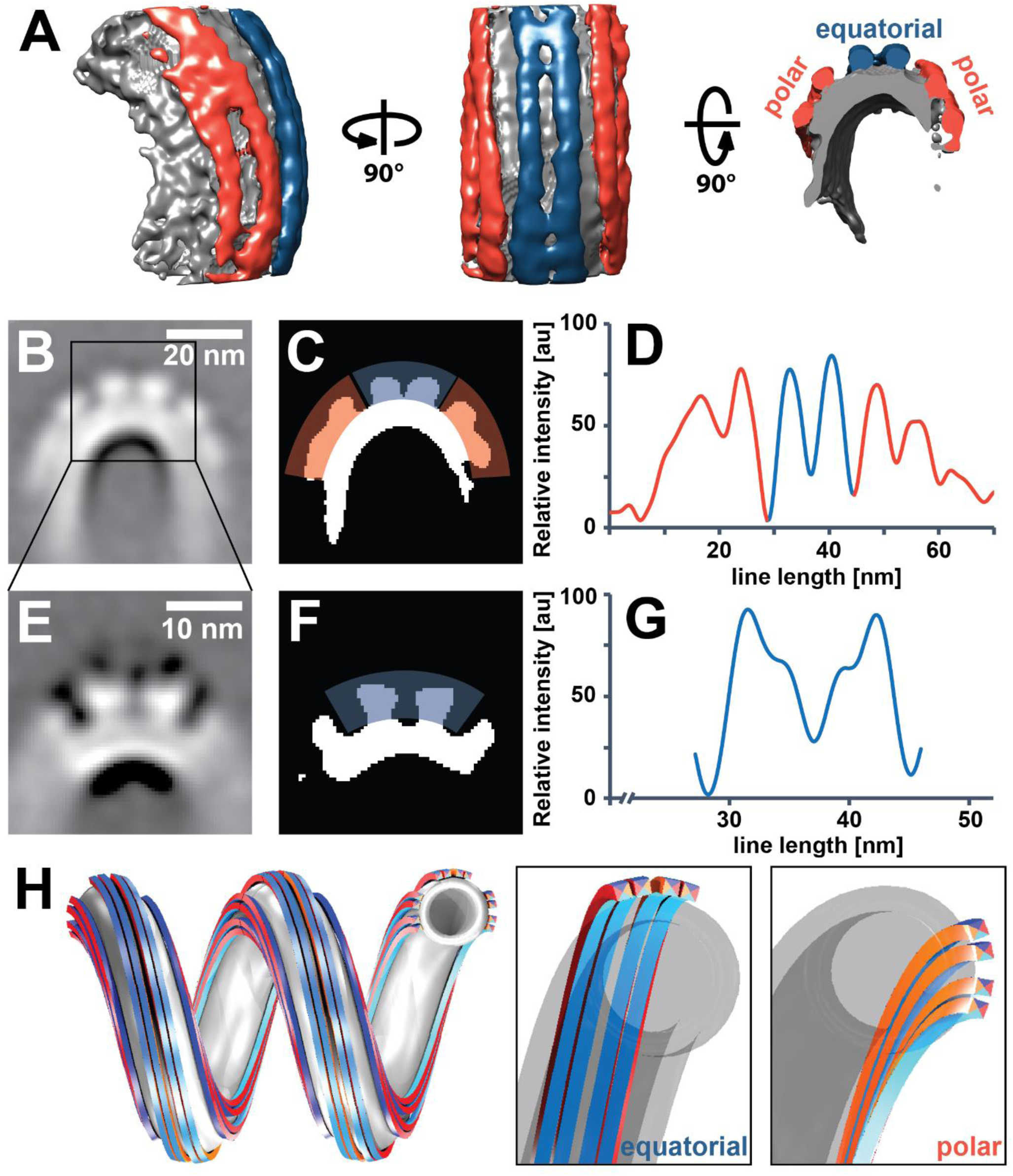
ESCRT-III filament bundles form distinct clusters on the surface of helical tubes. (A) Side view (left), top view (center) and cross-section (right) of a global subtomogram average showing filaments following the tube axis, in the equatorial (blue) and polar (red) binding mode, respectively. (B) Sum projection of a central segment of the tube in (A) showing filaments on the outer surface of the helical tube, organized as one equatorial and two polar clusters. Scale bar 20 nm. (C) Equatorial (blue) and polar (red) filament cluster highlighted on the thresholded image (B). (D) Intensity profile of protein density in (C). (E) Projection of a refined map of the equatorial cluster showing that both filaments of the cluster are made of two strands each. Scale bar 10 nm. (F) Thresholded image of (E). (G) Intensity profile of protein density in (E). (H) 3D model of one equatorial and two polar filament bundles, each formed from two double-strands, on a helical membrane tube (grey). All filaments are identical, except that equatorial and polar filaments bind the membrane through the cyan and orange interfaces, respectively (insets). Filaments in the two hemispheres are shown as antiparallel.

With further STA focused on the equatorial cluster, we reconstructed a map of this area (32 Å resolution), revealing that the two equatorial filaments contain two strands each (Fig. 2E-G, Fig. S2B). The filaments bundle in a plane parallel to the tube’s helical axis and their membrane binding area is on the bundle’s inside, also parallel to the helical axis, as observed in previously described ESCRT-III heteropolymers (*6*). Yet, in our case, both strands appear to be interacting with the membrane. The filaments in the polar clusters, based on their thickness, could be double-stranded as well, though our reconstructions were unable to resolve the substructure directly. In contrast to the equatorial filaments, however, the bundling plane of the polar filament strands is perpendicular to the helical axis, as is its membrane-binding interface (Fig. 2H). This orientation fits the double-stranded spirals formed by Snf7/Vps24/Vps2 on flat bilayers (*11*). Overall, the architectures of equatorial and polar filaments appear to be similar: both are composed of at least two double-stranded filaments, bundled together as a helical ribbon along the surface of the tube. However, the geometry of the helical tube makes it impossible that all filaments have the same path and bind the membrane with the same interface (Fig. 2H). For the same reasons, interactions between filaments within a bundle cannot be the same in polar filaments and equatorial filaments. While the possibility that ESCRT-III molecules bind their target membranes with two different orientations seems *a priori* unexpected, existing structural studies have reported different membrane binding interfaces for Snf7 versus CHMP1B (*6, 16*).

To clarify the interplay between the elasticity of the ESCRT-III filaments and that of the membrane in determining the shape of the helical tube, we sought to analyze the spontaneous shape of ESCRT-III filaments without a helical membrane tube for higher-resolution imaging. By growing Snf7/Vps24/Vps2 filaments in the presence of detergent-solubilized lipids, helical ribbons formed without membrane tubes during detergent removal (Fig. S3A-C). Most of these tube-less, helical ribbons assembled into sharp zigzag shapes (Fig. 3A, red arrows in Fig S3A-C), a smaller population appeared sinusoidal (Fig. 3B, blue arrows in Fig S3A-C), and a third ribbon population displayed significantly larger ribbons with varying strand numbers and diameters (Fig. 3C, yellow arrows in Fig. S3A-C). We used single-particle averaging approaches to analyze these tube-less helical protein filament ribbons and determined 2D class averages (Fig. 3D-F). The overall appearance of these sinusoidal ribbons suggests they comprise multi-stranded filaments that could orient along a helical path similar to that of the equatorial filaments we observe bound to the helical membrane tubes (Fig. 3E).

**Fig. 3:**
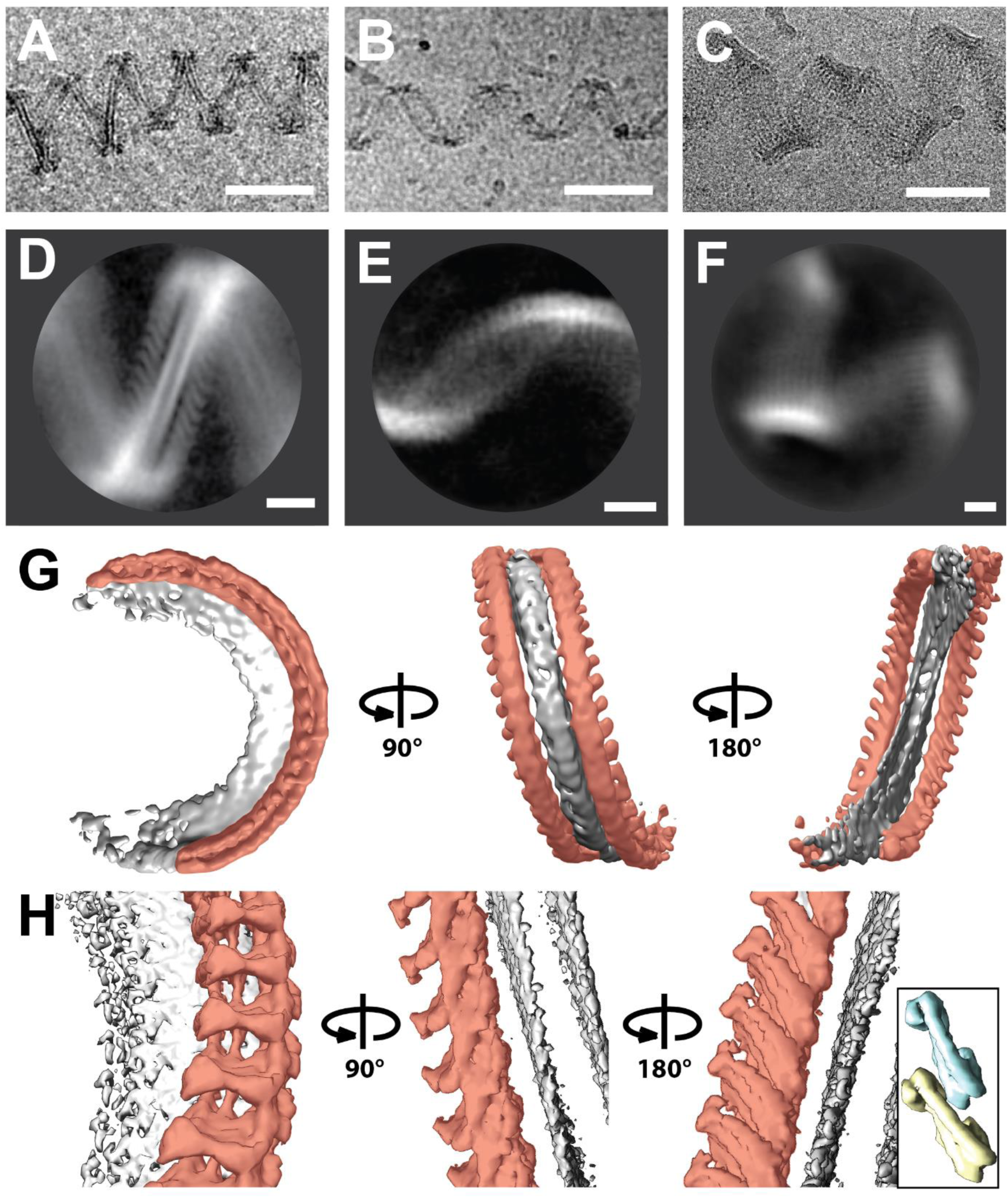
Organization of tube-less ESCRT-III filaments. Electron micrographs (A-C, scale bars 100 nm) and 2D class averages (D-F, scale bars 10 nm) showing different tube-less, helical ESCRT-III filament bundles formed upon detergent removal. The majority of ribbons adopted a zigzag shape (A, D), others appeared sinusoidal (B, E) and a third set consisted of helical ribbons with higher strand numbers (C, F). (G) Unmasked 3D average of (A, D) shows that the center of the ribbon is a helical bicelle with its plane perpendicular to the tube axis (grey). There are two anti-parallel double-stranded filaments on both sides of the bicelle (red). (H) 3D average as in (A, C) with an asymmetric mask that included only one double-stranded filament (red). Inset: scale-matched densities corresponding to two closed-conformation IST1 subunits from EMD-6461 (*6*) are shown for comparison.

Analysis of the more ordered “zigzag” structure (Fig. 3A, D) led to a 3D reconstruction at 15 Å resolution. This structure revealed a helical ramp formed around a membrane bicelle, a tension-less lipid bilayer stabilized by detergents, with the bicelle plane oriented perpendicular to the helix axis. On both sides of the bicelle, we observed filamentous polymers with dimensions consistent with other double-stranded ESCRT-III structures (*6, 10, 11*). Considering the apparent subunit tilt on both sides of the bicelle, these appear to be anti-parallel to each other (Fig. 3D, G).

We confirmed the anti-parallel orientation of the two polymers by a 3D reconstruction at a higher resolution (11 Å) that was computed by focusing on one side of the bicelle only (Fig. 3H). The subunits appear to polymerize like previously described ESCRT-III heteropolymers and were oriented along a similar helical path. Surprisingly, both strands appear to interact with the membrane, and their membrane-binding interface is oriented perpendicular to the main helical axis (Fig. 3H). The interface is therefore perpendicular to that postulated for CHMP1B, which is parallel to the helix axis (*6*). Molecular docking allows fitting both filaments with crystal structures of subunits in the open (*D. melanogaster* CHMP4B homolog Shrub, PDB 5J45 (*17*); yeast Snf7, PDB 5FD9 (*16*)) and closed conformation (Human CHMP3; PDB 3FRT (*18*)), respectively (Fig. S3D), with inter-subunit connectivity similar to known ESCRT-III heteropolymer structures (*6*). The resolution of the map, however, did not allow us to discern unambiguous conformations for the subunits of either strand. Nevertheless, the zigzag tube-less ribbon’s architecture is compatible with the polar filaments on helical tubes, and confirm that the polar filaments of the helical tube are also double-stranded. Our results confirm that ESCRT-III filaments can bind the membrane with two different orientations and form membrane structures with complex curvatures (Fig. 2H).

To understand how ESCRT-III filaments and the membrane influence each other’s conformation, we developed a mathematical model that describes the competition between filament and membrane rigidities, membrane tension and filament-membrane binding energy (Supplementary Information). In a first approach, we considered a helical scaffold of fixed geometry (consistent with the modest – less than 20% – deformation induced by the addition of membrane). Adding the membrane to this scaffold, we parametrized the energy difference between filament binding modes by an energy *μ* per unit filament length, where *μ* > 0 promotes polar filaments over equatorial ones and thus favors helical tubes (Fig. 4A). Indeed, straight tubes are coated only by equatorial filaments. We found that helical tubes are always favored at high membrane tension *σ*, and that lowering the tension leads to an increase of the membrane tube radius *r*, with different effects as a function of the value of *μ*. For high values of *μ* helical tubes remain stable at all *σ*. For lower values of *μ, r* increases significantly before reaching a *μ*-dependent critical value *r*_*c*_ where the system transitions to a straight tube (Fig. 4B). As a result, radii larger than *r*_*c*_ cannot occur, and the observation of relatively thick tubes with *r*_*exp*_ = 12.1 nm thus implies that 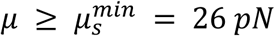 (Fig. 4A). This corresponds to a Snf7/Vps24/Vps2 binding energy difference of 2 *k*_*B*_*T* per monomer, a value compatible the previously estimated membrane-binding energy of Snf7 polymers alone (about 4 *k*_*B*_*T* per monomer (*5*)).

**Fig. 4:**
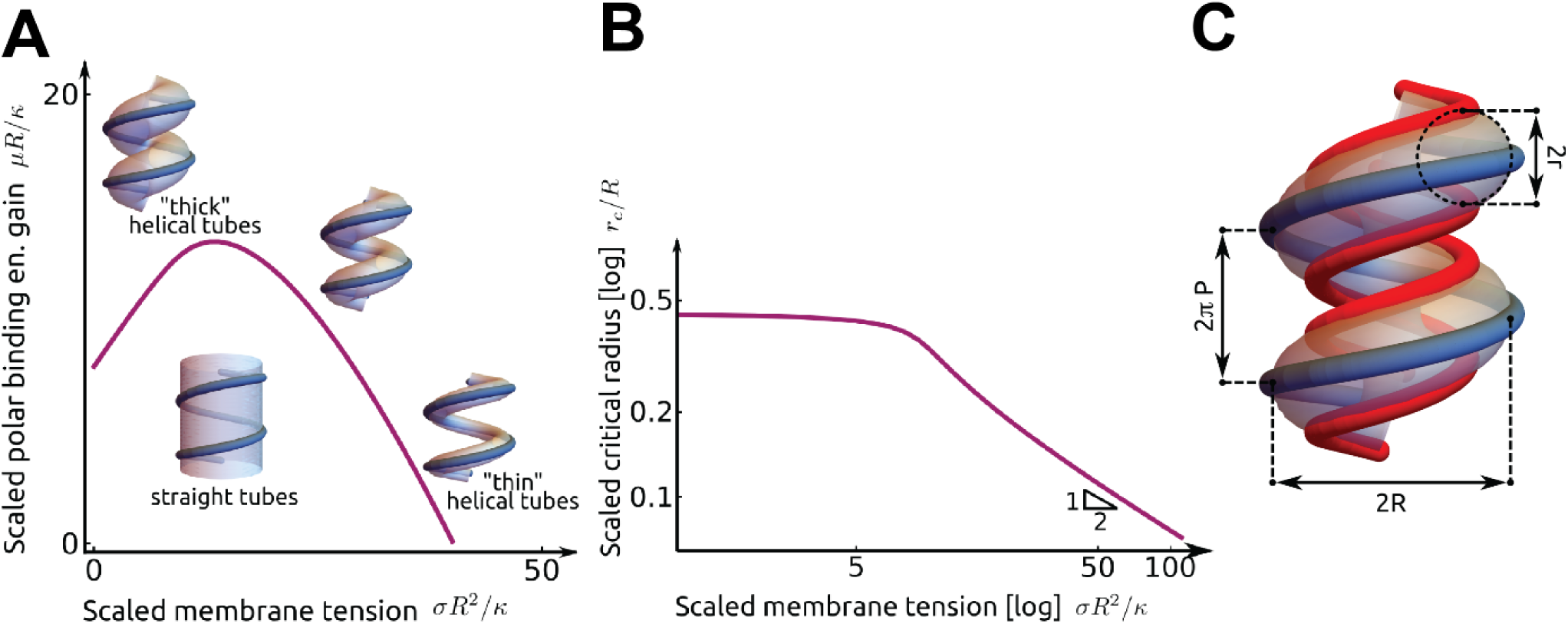
Mathematical modeling of helical tubes’ mechanical equilibrium. (A) Phase diagram showing the energetically favored shape between straight and helical tubes, as a function of energy gain μ associated with helical tubes with membrane tension σ. The solid purple line is the phase boundary. (B) Critical radius *r*_*c*_ of the tube at the transition from straight to helical as a function of surface tension *σ*. (C) Schematic of the more detailed filament elasticity model, which clusters together the filaments bound in the equatorial (blue) and polar modes (red).

We used the magnitude of the deformation of the Snf7/Vps24/Vps2 filaments in the presence of the membrane to infer their mechanical rigidities as well as *μ*. Using the more detailed filament geometry (Fig. 4C), we endowed the Snf7/Vps24/Vps2 filaments with bending and torsional rigidities, with the former being characterized by the filament persistence length lp. We found that the differences in filament radii and pitch observed as a function of the presence (Fig. 2) or absence (Fig. 3) of membrane imply that 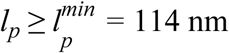, and *μ ≥* 5 *k*_*B*_*T* /monomer, slightly larger than the lower bound on *μ* derived above, implying that helical tubes are more favorable than straight ones in the whole range of predicted rigidities. Another possible estimate was obtained by setting 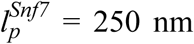 (*5*) for Snf7-only filaments. This implies a torsional stiffness of 45 nm *k*_*B*_*T,* comparable to that of DNA at low tension (*19*), as well as a differential binding energy of *μ* = 15 *k*_*B*_*T*/monomer, suggesting that Vps24 and Vps2 may be significant contributors of the binding of ESCRT-III filaments to lipid membranes.

Our findings support the hypothesis that the assembly of multiple strands of ESCRT-III triggers a buckling transition. Increasing filament torsional rigidity, in addition to bending rigidity (*5*), could trigger the buckling transition. In this case, the filament in the flat spiral would be pre-constrained (no torsion), and the increase of its torsional rigidity when it pairs with additional strands would allow the new composite filament to adopt a conformation closer to its preferred torsion (helical). Under these circumstances, a buckling transition could be possible with a lower number of ESCRT-III subunits, with compositional heterogeneity, explaining why our previous model (*5*) requires more subunits than are found at sites of intraluminal vesicle formation (*20*).

Considering the helical path of ESCRT-III assemblies, structural studies have identified several membrane-interacting surfaces on the inside (*6*) and the outside (*16*) of the helix, and we identify here a third surface perpendicular to those. This may reveal a more complex picture of the filament shape transition involved in membrane deformation. If ESCRT-III subunits change their membrane-binding interface during their deformation, this could allow a filament to roll on the membrane and generate torque along the filament axis as another source of membrane strain.

Shape buckling and torque may originate from subunits from the leading strand being exchanged by different subunits that bind the membrane with a different preferred orientation. We have shown that subunit turnover, and incorporation of different subunits are both necessary for ESCRT-III-mediated membrane (*11, 21*). Additionally, or alternatively, the formation of a secondary membrane-binding filament parallel to the leading strand (*11*) could change the membrane-binding interface orientation, forcing the membrane to adopt a tubular shape. Unfortunately, our data did not allow us to establish whether polar and equatorial binding modes reflect different heteropolymer stoichiometries or different conformations of the same proteins forming the heteropolymer.

Whereas bending and torsional rigidity define the helical nature of our tube-less ESCRT-III filaments, the shape of the helical membrane tube is inherently different. The helical tube is, therefore, the result of a competition between the scaffold shape and the membrane properties, particularly membrane tension. Though membrane forces probably also affect the geometries of other protein scaffolds, we show here a first semi-rigid scaffold whose geometry is significantly affected by membrane properties.

## Supporting information

Supplementary mathematical modeling

## Acknowledgments

The authors would like to thank Alexander Myasnikov, Arthur Melo and Wim Hagen for help with electron microscopy data collection and processing.

## Author contributions

Conception and design: JMvF and AR; data acquisition, analysis and interpretation: JMvF, LB, NT, IJ, AF, ML and AR; theoretical model: LB and ML; writing (original draft): JMvF, LB, ML and AR; writing (review and editing): JMvF, LB, NT, IJ, AF, ML and AR

## Data and materials availability

All data needed to evaluate the conclusions in this paper are presented here or in the supplementary materials. Structural data is available from the Electron Microscopy Data Bank, accession numbers for electron density maps are EMD-10136, EMD-10137, EMD-10138 and EMD-10138.

## Supplementary Materials

Materials and Methods

Supplementary Mathematical Modeling

Figures S1-S3

Movies S1-S2.

