## Supplementary mathematical modeling for "Anisotropic ESCRT-III architecture governs helical membrane tube formation"

### Anisotropic ESCRT-III architecture governs helical membrane tube formation
# —
### Supplementary mathematical modeling

Moser Von Filseck *et al.*

Sec. (1) of this supplement demonstrates that helical tubes can be more stable than straight ones, and uses the observations of the main text to infer a bound on the energy difference between the two filament binding modes described there. In Sec. (2), we propose a model of helical tube stabilized by a deformable scaffold of Snf7/Vps24/Vps2 heteropolymers. Force-balance conditions allow us to estimate the bending and torsional rigidities of Snf7/Vps24/Vps2 heteropolymers, as well as the binding energy difference between polar and equatorial filaments. Finally, in Sec. (3) we present a detailed derivation of differential geometry results used in the other sections. Most of the results are discussed in the main text.

#### 1 Helical tubes can be more stable than straight tubes

In this section, we compare the stability of the helical and straight tubes illustrated in Fig. (4A) of the main text. We show that helical tubes are more favorable than straight ones for high enough membrane tensions, and that the radius of the tube at the transition depends on the difference in membrane binding energy between polar and equatorial filaments [respectively illustrated in blue and red in Fig. (SM1)]. We approximate the membrane scaffolding function of the filaments by a single, effective outer helix that fixes the outer membrane radius [Fig. (4A), main text]. We moreover assume that this scaffold is undeformable, consistent with the observation made in the main text that its dimensions change only by a modest amount (less than $20\%$) when membrane is added to it.

Our description includes two free energy contributions, from membrane elasticity and from filament-membrane binding. We compute the former by modeling the membrane as a thin liquid sheet governed by the Helfrich free energy [3]

$$\mathcal{F}_{\text{Helfrich}} = \int_{\mathcal{A}} \mathrm{d}\mathcal{A} \left[\frac{\kappa}{2}(2H)^2 + \sigma\right], \tag{1}$$

where the integral runs over the area of the membrane, $\kappa \simeq 20\ k_B T$ is its bending rigidity, $H$ its mean curvature and $\sigma$ its surface tension. Now turning to the filament-membrane binding contribution to the energy, we attribute different energies to the polar and equatorial filament binding modes. While filaments can only bind to straight tubes through the equatorial mode, helical tubes are covered by both polar and equatorial filaments, implying an energy difference $-2/3 \times \mu$ between the filament binding energy per unit length in the helical tube compared to straight tubes (*i.e.*, $\mu > 0$ favors helical tubes). By the factor $2/3$, we account for the fraction of scaffolding filaments binding in the polar mode, consistently with the experiments. The differential binding energy per monomer can be written as $\mu \times 1/8 \times 3\,\mathrm{nm}$, where the factor $1/8$

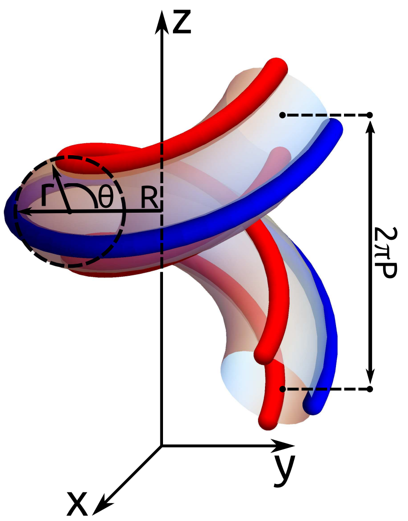


Figure SM1: Schematic of the deformable polymer helix model. A scaffold made of three effective filaments stabilizes a helical tube. Each effective filament accounts for two sets of two Snf7/Vps24/Vps2 sub-filaments, consistent with the average number of filaments observed around helical tubes in the experiments [Fig. (2H), main text]. Blue filaments are bound in the "equatorial" binding mode, while red ones are bound in the "polar" mode. The two polar filaments are localized at $\theta = \pm\pi/2$, while the equatorial filament is localized at $\theta = \pi$.

accounts for the four sets of two polar Snf7/Vps24/Vps2 sub-filaments included in our model helix [Fig. (2H), main text], and 3 nm is the typical monomer spacing.

We now compare the free energy of straight and helical tubes for a fixed total length $\mathcal{L}$ of helical scaffold of radius $R$ and pitch $2\pi P$ [Fig. (4C), main text]. Approximating the straight tube to a cylinder, Eq. (1) reduces to

$$\frac{\mathcal{F}_{\text{straight}}}{\mathcal{L}} = \frac{2\pi RP}{\left(R^2+P^2\right)^{1/2}}\left(\frac{\kappa}{2}\frac{1}{R^2}+\sigma\right). \tag{2}$$

Now considering the helical tube, we model the membrane as a tube of constant radius $r$ with a helical center line. This geometry implies expressions of the tube curvature and area detailed in Sec. (3), yielding the free energy

$$\frac{\mathcal{F}_{\text{helical}}(r)}{\mathcal{L}} = 2\pi\left[\frac{P^2+(R-r)^2}{P^2+R^2}\right]^{1/2}\left\{\frac{\kappa}{2r}\left[1-\left(\frac{r(R-r)}{P^2+(R-r)^2}\right)^2\right]^{-1/2}+r\sigma\right\}-\frac{2}{3}\mu, \tag{3}$$

where the additional $\mu$ term accounts for the lower binding energy of the helical tube. Using $R = 41.9\,\text{nm}$ and $P = 8.5\,\text{nm}$, we numerically minimize $\mathcal{F}_{\text{helical}}$ over $r$ and compare it to $\mathcal{F}_{\text{straight}}$. The parameter regions where either type of tubes is more stable are illustrated in Fig. (4A) of the main text.

#### 2 Inferring filament rigidities from helical tube geometries

Compared to an isolated Snf7/Vps24/Vps2 helix, adding a membrane of known rigidity deforms the polymer into a larger helix, as described in the main text. In this section, we use the magnitude of this deformation to infer the bending and torsional rigidities of the helix as well as the difference in membrane binding energy between the polar and equatorial filaments.

The model used here is more detailed than the one of Sec. (1), and takes into account the flexibility of the filaments. In the model, a helical membrane tube is bound to two polar (red) and one equatorial (blue) filaments [Fig. (SM1)]. Every model filament accounts for two Snf7/Vps24/Vps2 filaments, each of which is made of two sub-filaments, as reported in Fig. (2H) of the main text. The difference in binding energy per unit length between the two types of filaments is denoted by $\mu$ as in Sec. (1). Denoting the radius of the blue polymer by $R$ and its pitch by $2\pi P$, the total length of red filaments is $\mathcal{L}^R = 4\pi n\sqrt{(R-r)^2 + P^2}$ and the length of the blue filaments is $\mathcal{L}^B = 2\pi n\sqrt{R^2 + P^2}$, where $n$ denotes the number of turns around the vertical axis. We consider the most energetically favorable configuration of the system at fixed total polymer length $\mathcal{L} = \mathcal{L}^R + \mathcal{L}^B$ and membrane area $\mathcal{A}$. Denoting by $a = \mathcal{A}/\mathcal{L}$ the membrane surface area per unit polymer length, we use the expression of $\mathcal{A}$ given in Sec. (3) to express the pitch $2\pi P$ as

$$P = \left[\frac{(2 - 2\pi r/a)^2 (R-r)^2 - R^2}{1 - (2 - 2\pi r/a)^2}\right]^{1/2}, \tag{4}$$

where $r$ is the radius of the membrane tube. The free energy of the system is the sum of a membrane and a polymer contribution: $\mathcal{F} = \mathcal{F}_{\text{membrane}} + \mathcal{F}_{\text{polymer}}$. The membrane free energy is given by Eq. (1), noting that the surface tension term only contributes a (physically irrelevant) constant to the free energy due to the constraint of fixed $\mathcal{A}$. The polymer free energy is given by:

$$\frac{\mathcal{F}_{\text{polymer}}}{\mathcal{L}} = \ell^R\left[\frac{k_b}{2}\left(c^R - c_0^R\right)^2 + \frac{k_\tau}{2}\left(\tau^R - \tau_0^R\right)^2 - \frac{\mu}{2}\right] + \ell^B\left[\frac{k_b/4}{2}\left(c^B - c_0^B\right)^2 + \frac{k_\tau}{2}\left(\tau^B - \tau_0^B\right)^2\right], \tag{5}$$

where the superscripts $R$ and $B$ denote quantities related to red and blue filaments, respectively. Here $\ell^R = \mathcal{L}^R/\mathcal{L}$, $\ell^B = \mathcal{L}^B/\mathcal{L}$ and $c_0 = R_0/(R_0^2 + P_0^2)$ and $\tau_0 = P_0/(R_0^2 + P_0^2)$ denote the spontaneous curvature and torsion of the filaments, and are given by the radius $R_0$ and pitch $2\pi P_0$ of the tube-less helices. The values of the curvature and torsion of the deformed filaments are given as

$$c^R = \frac{R-r}{(R-r)^2 + P^2}, \quad c^B = \frac{R}{R^2 + P^2}, \quad \tau^R = \frac{P}{(R-r)^2 + P^2} \quad \text{and} \quad \tau^B = \frac{P}{R^2 + P^2}. \tag{6}$$

The differential binding energy per unit length $\mu$ is rescaled by $1/2$, so that its definition is the same as in Sec. (1), where one model filament accounted for four sets of two polar Snf7/Vps24/Vps2 sub-filaments, whereas here one model filament accounts for two sets of two sub-filaments. While both types of filaments have the same torsional stiffness $k_t$, in Eq. (5) the bending stiffness $k_b$ of the red filaments is four times larger than that of the blue. Indeed, as we show in the main text, both red and blue filaments consist of two parallel sub-filaments of comparable thickness, but while red filaments bend along the sub-filaments' binding direction, blue filaments bend along the orthogonal direction. Just like it is much easier to bend a piece of cooked tagliatelle pasta in its thin than its thick direction, bending the red filaments is thus more costly than bending the blue ones. To express this notion quantitatively, we approximate each set of two filaments as an elastic rod with a rectangular cross-section of aspect ratio equal to 2. Applying the classical result of Ref. [4] this implies a ratio of bending stiffnesses of $2^2 = 4$.

We next express the condition of mechanical equilibrium of our system, which relates the helix' mechanical parameters with its observed dimensions. We thus insert Eqs. (4) and (6) into Eq. (5) to express the free energy of our system as a function of $R$ and $r$ only, then write the two force balance equations $\partial\mathcal{F}/\partial r = 0$ and $\partial\mathcal{F}/\partial R = 0$. Finally, we insert the numerical values of $R = 41.9\,\text{nm}$, $r = 12.1\,\text{nm}$, $R_0^B = 17.1\,\text{nm}$, $P_0^B = 8.9\,\text{nm}$, $R_0^R = 23.4\,\text{nm}$ and $P_0^R = 6.6\,\text{nm}$ reported in the main text, as well as $a = 22.7\,\text{nm}$ obtained from Eq. (15) using $P = 8.5\,\text{nm}$. This results in a set of two equations relating the unknown parameters $k_b$, $k_t$ and $\mu$, which can be recast as:

$$\frac{k_b}{\kappa/c_0^R} = -1.14 + 0.30 \times \frac{\mu}{\kappa c_0^R} \tag{7a}$$

$$\frac{k_t}{\kappa/c_0^R} = -0.32 + 0.02 \times \frac{\mu}{\kappa c_0^R}. \tag{7b}$$

Since the mechanical stability of our helix requires that $k_t \geq 0$, Eqs. (7) further imply that $\mu > \mu_k^{\text{min}} \simeq 52\,\text{pN}$, corresponding to a differential binding energy per monomer bigger than $\mu_k^{\text{min}} \times 1/8 \times 3\,\text{nm} \simeq 5\,k_BT$, and $k_b > k_b^{\text{min}} \simeq 8 \times 10^{-27}\,\text{J}\cdot\text{m}$.

The value of $k_b^{\text{min}}$ sets a lower bound $\ell_p^{\text{min}}$ for the persistence length $\ell_p$ of individual Snf7/Vps24/Vps2 sub-filaments. To derive this bound, we note that our model filaments account for two sets of two Snf7/Vps24/Vps2 sub-filaments. We assume that each set of two sub-filaments responds to bending like an isotropic rod of rectangular section, with short side $d$ and long side $2d$. We further assume that sub-filaments respond to bending like isotropic rods of square section with side $d$. This implies

$$\ell_p^{\text{min}} = \frac{1}{2} \times \frac{d^4/12}{(2d^3)\,d/12} \times \frac{k_b^{\text{min}}}{k_BT} \simeq 114\,\text{nm}, \tag{8}$$

on the basis of the classical theory of elasticity [4].

We conclude this section by estimating the actual torsional stiffness of Snf7/Vps24/Vps2 sub-filaments, as well as their differential binding energy per monomer, assuming that the actual persistence length of Snf7/Vps24/Vps2 sub-filaments is that of Snf7, $\ell_p^{\text{Snf7}} = 250\,\text{nm}$ [1]. The differential binding energy per monomer is $\mu^* \times 1/8 \times 3\,\text{nm} \simeq 15\,k_BT$, and $\mu^*$ is the solution of Eq. (7a) for $k_b = \ell_p^{\text{Snf7}} \times k_BT$. Let us call $k_t^*$ the solution of Eq. (7b) for $\mu = \mu^*$. Once again, we assume that each set of two sub-filaments and sub-filaments respond to torsion like elastic rods, with rectangular and square section, respectively. Then, since each model filament accounts for two sets of two Snf7/Vps24/Vps2 sub-filaments, classical elasticity [4] implies that the torsional persistence length of a sub-filament is

$$\frac{1}{2} \times \frac{\beta_1 d^4}{\beta_2\,(2d)\,d^3} \times \frac{k_t^*}{k_BT} \simeq 45\,\text{nm}, \tag{9}$$

where $\beta_1 = 0.141$ and $\beta_2 = 0.229$ are geometrical constants available in Ref. [2].

### 3 Differential geometry of helical tubes

Here we compute the area and bending energy of a helical tube. To perform this calculation, we compute the first $(g)$ and second $(b)$ fundamental forms of the surface and use them to express its local differential area $\mathrm{d}\mathcal{A}$ and mean curvature $H$.

We first parametrize the centerline of the helical tube. This centerline is a regular helix with radius $R - r$ and pitch $2\pi P$. Denoting by $s \in [0, \mathcal{S} = 2\pi n\sqrt{(R-r)^2 + P^2}]$ its arc length, where $n$ is the number of turns around the z-axis, its position vector is

$$\mathbf{h}(s) = R\left[\cos\left(\frac{s}{\sqrt{(R-r)^2 + P^2}}\right)\widehat{\mathbf{x}} + \sin\left(\frac{s}{\sqrt{(R-r)^2 + P^2}}\right)\widehat{\mathbf{y}}\right] + \frac{sP}{\sqrt{(R-r)^2 + P^2}}\,\widehat{\mathbf{z}}. \tag{10}$$

To parametrize the tubular surface, we take advantage of the Frenet – Serret frame of the curve $\mathbf{h}(s)$, generated by its unit tangent $\widehat{\mathbf{t}}$, normal $\widehat{\mathbf{n}}$ and binormal $\widehat{\mathbf{b}}$

$$\widehat{\mathbf{t}}(s) = \frac{\partial \mathbf{h}}{\partial s}; \qquad \widehat{\mathbf{n}}(s) = \frac{(R-r)^2 + P^2}{R-r}\frac{\partial^2 \mathbf{h}}{\partial s^2}; \qquad \widehat{\mathbf{b}}(s) = \widehat{\mathbf{t}} \times \widehat{\mathbf{n}}, \tag{11}$$

yielding the position vector of the surface as

$$\boldsymbol{\Sigma}(\theta, s) = \mathbf{h}(s) + r\left(\cos\theta\,\widehat{\mathbf{n}}(s) + \sin\theta\,\widehat{\mathbf{b}}(s)\right), \tag{12}$$

where $r$ is the membrane radius and $\theta \in [0, 2\pi]$. The covariant metric tensor $(g_{\alpha\beta})$ associated to the membrane is:

$$(g_{\alpha\beta}) \doteq \begin{pmatrix} \frac{\partial \boldsymbol{\Sigma}}{\partial \phi}\cdot\frac{\partial \boldsymbol{\Sigma}}{\partial \phi} & \frac{\partial \boldsymbol{\Sigma}}{\partial \theta}\cdot\frac{\partial \boldsymbol{\Sigma}}{\partial \phi} \\ \frac{\partial \boldsymbol{\Sigma}}{\partial \theta}\cdot\frac{\partial \boldsymbol{\Sigma}}{\partial \phi} & \frac{\partial \boldsymbol{\Sigma}}{\partial \theta}\cdot\frac{\partial \boldsymbol{\Sigma}}{\partial \theta} \end{pmatrix}, \tag{13}$$

where the indices of $g$ can be either $\phi$ or $\theta$. The differential area of the tubular surface reads

$$\mathrm{d}\mathcal{A} = \sqrt{g}\,\mathrm{d}\theta\,\mathrm{d}s = r\left[1 - \frac{r(R-r)\cos\theta}{(R-r)^2 + P^2}\right]\mathrm{d}\theta\,\mathrm{d}s \tag{14}$$

where $g$ is the determinant of the metric tensor. This yields the total area for the tubular membrane:

$$\mathcal{A} = 2\pi r \mathcal{S}. \tag{15}$$

The second fundamental form $(b_{\alpha\beta})$ associated to the surface $\boldsymbol{\Sigma}$ reads in covariant representation

$$(b_{\alpha\beta}) \doteq \begin{pmatrix} \frac{\partial^2 \boldsymbol{\Sigma}}{\partial \phi^2}\cdot\widehat{\mathbf{N}} & \frac{\partial^2 \boldsymbol{\Sigma}}{\partial \theta \partial \phi}\cdot\widehat{\mathbf{N}} \\ \frac{\partial^2 \boldsymbol{\Sigma}}{\partial \theta \partial \phi}\cdot\widehat{\mathbf{N}} & \frac{\partial^2 \boldsymbol{\Sigma}}{\partial \theta^2}\cdot\widehat{\mathbf{N}} \end{pmatrix}, \tag{16}$$

where the unit vector $\widehat{\mathbf{N}}$ normal to $\boldsymbol{\Sigma}$ is defined as:

$$\widehat{\mathbf{N}}(\theta, s) = \frac{\frac{\partial \boldsymbol{\Sigma}}{\partial \theta} \times \frac{\partial \boldsymbol{\Sigma}}{\partial \phi}}{\|\frac{\partial \boldsymbol{\Sigma}}{\partial \theta} \times \frac{\partial \boldsymbol{\Sigma}}{\partial \phi}\|}. \tag{17}$$

The mean curvature is obtained by tracing the second fundamental form

$$2H = g^{\alpha\beta} b_{\alpha\beta} = -\frac{P^2 + R^2 - 2rR\cos\theta}{r\left(P^2 + R^2 - rR\cos\theta\right)}, \tag{18}$$

where $\left(g^{\alpha\beta}\right) = (g_{\alpha\beta})^{-1}$ is the contravariant representation of the metric tensor and summation over repeated indices is implied. Finally, we combine Eqs. (3) and (18) to compute the integral appearing in the bending energy of a helical membrane tube:

$$\int_{\boldsymbol{\Sigma}} \mathrm{d}\mathcal{A}\,(2H)^2 = \int_0^{\mathcal{S}} \mathrm{d}s \int_0^{2\pi} \mathrm{d}\theta\,\sqrt{g}\,(2H)^2 = \frac{2\pi\mathcal{S}}{r}\left\{1 - \frac{r^2(R-r)^2}{\left[(R-r)^2 + P^2\right]^2}\right\}^{-1/2}. \tag{19}$$

[4] L. D. Landau and E. M. Lifshitz. *Theory of Elasticity*.
