## Supplementary information for "Anisotropic ESCRT-III architecture governs helical membrane tube formation"

### Anisotropic bending stiffnesses of ESCRT-III heteropolymers dictate membrane curvature acquisition

**This PDF file includes:**

Materials and Methods
Figs. S1 to S3
Captions for Movies S1 to S2

**Other Supplementary Materials for this manuscript include the following:**

Movies S1 to S2
Supplementary mathematical modeling

**Materials and Methods**

Protein expression and purification

Proteins were expressed from plasmids encoding budding yeast Snf7 (Addgene no. 21492), Vps2 (Addgene no. 21494) and Vps24 (gift from James Hurley), and were purified as previously described (*1*).

Liposome preparation

1,2-dioleoyl-sn-glycero-3-phosphocholine (DOPC) and 1,2-dioleoyl-sn-glycero-3-phospho-L-serine (sodium salt) (DOPS) were purchased in solution from Avanti Polar Lipids and mixed at the desired molar ratio in chloroform. The lipid mix was dried first under a nitrogen stream and then under vacuum at 30 °C for 1 h before hydration with 100 mM NaCl 20 mM Hepes pH = 7.5. We made large unilamellar vesicles (LUVs) by extrusion of the hydrated lipid films using a Mini Extruder (Avanti Polar Lipids) and polycarbonate filters of pore size 0.2 µm (Whatman).

Formation of helical membrane tubes

At 4 °C, in 100 mM NaCl, 20 mM Hepes pH = 7.5, extruded LUVs made from DOPC/DOPS (60/40 mol/mol) (10 mM final) were incubated with 10 µM Snf7 for one hour, then Vps2 and Vps24 (5 µM each) were added and incubated overnight. For Cryo-EM, 4 µL of the sample were deposited on glow-discharged Quantifoil R2/2 200 mesh copper grids and plunge frozen in liquid ethane after a two-sided blot using a FEI Vitrobot. For Cryo-ET, we added 10 nm BSA-nanogold (Aurion) to the reaction prior to vitrification. For negative stain EM, the sample was diluted 1/10 in 100 mM NaCl, 20 mM Hepes pH = 7.5 before staining for 30 s with 2% uranyl acetate.

Formation of protein polymers on bicelles

We prepared micelles by solubilizing a dried lipid film made from DOPC/DOPS (60/40 mol/mol) at 25 °C in 100 mM NaCl, 20 mM Hepes pH = 7.5, 20 mM CHAPS (3-[(3-Cholamidopropyl)-dimethyl-ammonio]-1-propanesulfonate hydrate, Sigma-Aldrich) at a total lipid concentration of 12 mM. The following protocol is adapted from (*2*). In brief, micelles were homogenized by bath sonication and stirring at 25 °C for 1 h before addition of 4 µM Snf7, 2 µM Vps24 and 2 µM Vps2, making sure that the detergent concentration was above its critical micellar concentration after addition of all proteins. The sample was then gradually diluted four-fold over 30 min under agitation at 25 °C and further incubated for 5 h. For Cryo-EM, 4 µL of the sample were deposited on glow-discharged Quantifoil R1.2/1.3 300 mesh copper grids and plunge frozen in liquid ethane after a two-sided blot using a FEI Vitrobot.

Low-resolution electron microscopy data collection

Transmission electron micrographs of negatively stained liposomes and helical tubes were acquired on a FEI Tecnai G2 Sphera $LaB_6$ at 200 kV using a 4k x 4k FEI Eagle Camera. Vitrified liposomes and helical tubes were imaged in low-dose mode on the same instrument.

High-resolution cryogenic electron microscopy data collection

962 Transmission electron cryo-micrographs of helical filaments on bicelles were collected on a FEI Titan Krios XFEG microscope at the University of California San Francisco, USA, equipped with a GIF K2 Quantum System (Gatan) and operated by Serial-EM software at 300 kV. Micrographs were collected at a nominal magnification of 105,000 x in super-resolution mode, corresponding to a super-resolution pixel size of 0.69 Å. Images were collected as dose-fractionated stacks for a total of 80 frames (0.2 sec/frame) and total dose of 67.2 electrons/Å$^2$. Coma-free beam alignments were performed prior to data collection and automated data collection was conducted with SerialEM (*3*).

Electron Cryogenic Microscopy Data Processing

Each dose-fractionated image stack was processed using MotionCorr2 and binned by two to yield motion- and dose-weighted images with a pixel size of 1.38 Å/pixel (*4*). For the double-filament structures, helical filaments were picked manually using RELION 3.0, segmented into ~90% overlapping segments and additionally binned by two during extraction with a final pixel size of 2.76 Å/pixel (bin4). After rejecting unalignable segments during 2D classification, 35,087 single particle images were processed by 3D classification and 3D auto-refine with helical priors, but without imposing helical symmetry (*5, 6*). For the single-filament structure, a soft mask was employed to remove one filament from the model and all of the images were re-processed by 3D classification and 3D auto-refine with helical priors and helical symmetry, using RELION 3.0 software (twist rotation angle (degrees) = 6.7, rise along axis = 10.6 Å). Independent half maps were post-processed using automated procedures. Reported resolutions based on FSC 0.143 criteria (*7*).

Electron cryo-tomography data collection

73 tilt series were collected on a FEI Titan Krios XFEG microscope at the European Molecular Biology Laboratory, Heidelberg, Germany, equipped with a GIF K2 Quantum System (Gatan) and operated by Serial-EM software at 300 kV. The tilt series were collected using a dose-symmetric scheme ranging ± 61° with 2° increments and defoci between -2.5 µm and -3.5 µm. The nominal magnification was 65,000 x with a calibrated pixel size of 2.14 Å. Images were recorded in counting mode with five frames per tilt angle and a total dose of 120 $e^{-}$ $Å^{-2}$ (2 $e^{-}$ $Å^{-2}$ $s^{-1}$ per tilt angle).

Tomogram reconstruction and subtomogram averaging

Tilt series of combined frames were aligned using the gold fiducials markers in IMOD (*8*). Tomograms were then reconstructed from these aligned tilt series using weighted back-projection in IMOD. Tomograms were binned four times and a 3D Gaussian filter of radius 2 was applied to increase contrast. Tomograms were then filtered using Hide Dust in UCSF Chimera (*9*), or manually segmented and analysed using 3dmod from the IMOD suite (*8*).

We used the Dynamo software for particle extraction and subtomogram averaging (*10*). We selected 8150 particle positions along the centre of a helical axis of membrane tubes in 17 bin2 tomograms. The table containing these positions was used to extract $160^3$ pixel particles from bin2 tomograms (bin2 particles). Bin2 particles were used to compute a first reference-free subtomogram average using a single, manually selected, blurred particle as alignment template, yielding a map at 31.9 Å resolution. This first subtomogram average was then used as a template for the alignment of 2037 bin2 particles with a soft elliptical alignment and classification mask focusing on the equatorial filament cluster, yielding a map with a resolution 32.4 Å. Reported resolutions based on FSC 0.143 criterion (*7*)

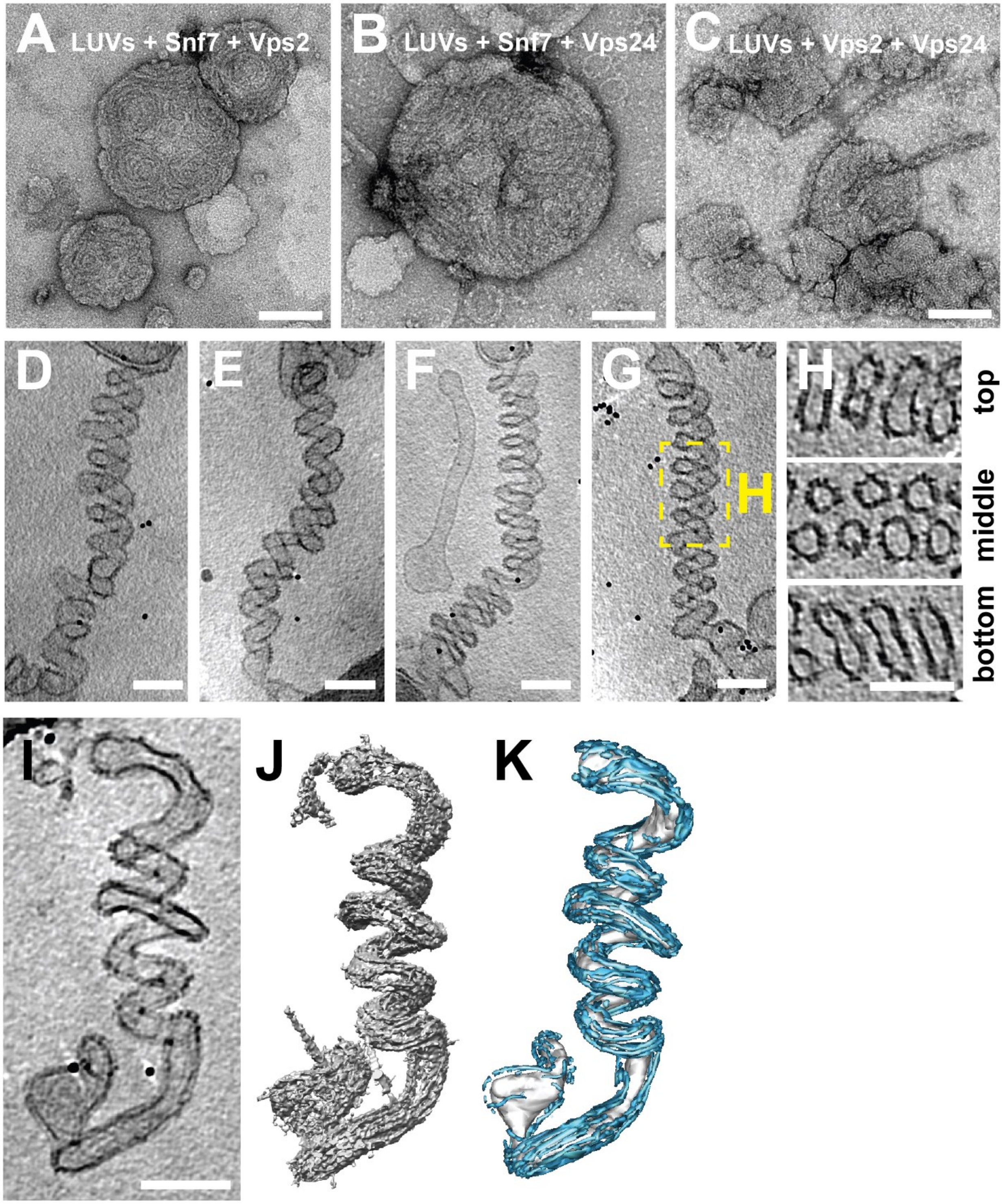
A LUVs + Snf7 + Vps2
B LUVs + Snf7 + Vps24
C LUVs + Vps2 + Vps24
D
E
F
G
H
H
top
middle
bottom
I
J
K

**Fig. S1: Helical tubulation of liposomes by ESCRT-III heteropolymers.** Representative electron micrographs showing undeformed liposomes decorated with Snf7 and Vps24 (A), Snf7 and Vps2 (B), and Vps24 and Vps2 (C) as observed on negatively stained samples. (D-G) Reconstructed Cryo-ET volumes of helical membrane tubes projected in Z displaying the variability in the tubes' helical parameters. (H) Slices through the top, centre and bottom of the helical tube shown in (G) illustrating its tubular-helical shape. Further example of reconstructed Cryo-ET volume of helical membrane tubes projected (I) and after filtering (J) or manual segmentation (K) showing the organisation of protein filaments along the tube axis. In (K), densities attributed to membrane and protein filaments are depicted in grey and cyan, respectively. All scale bars 100 nm.

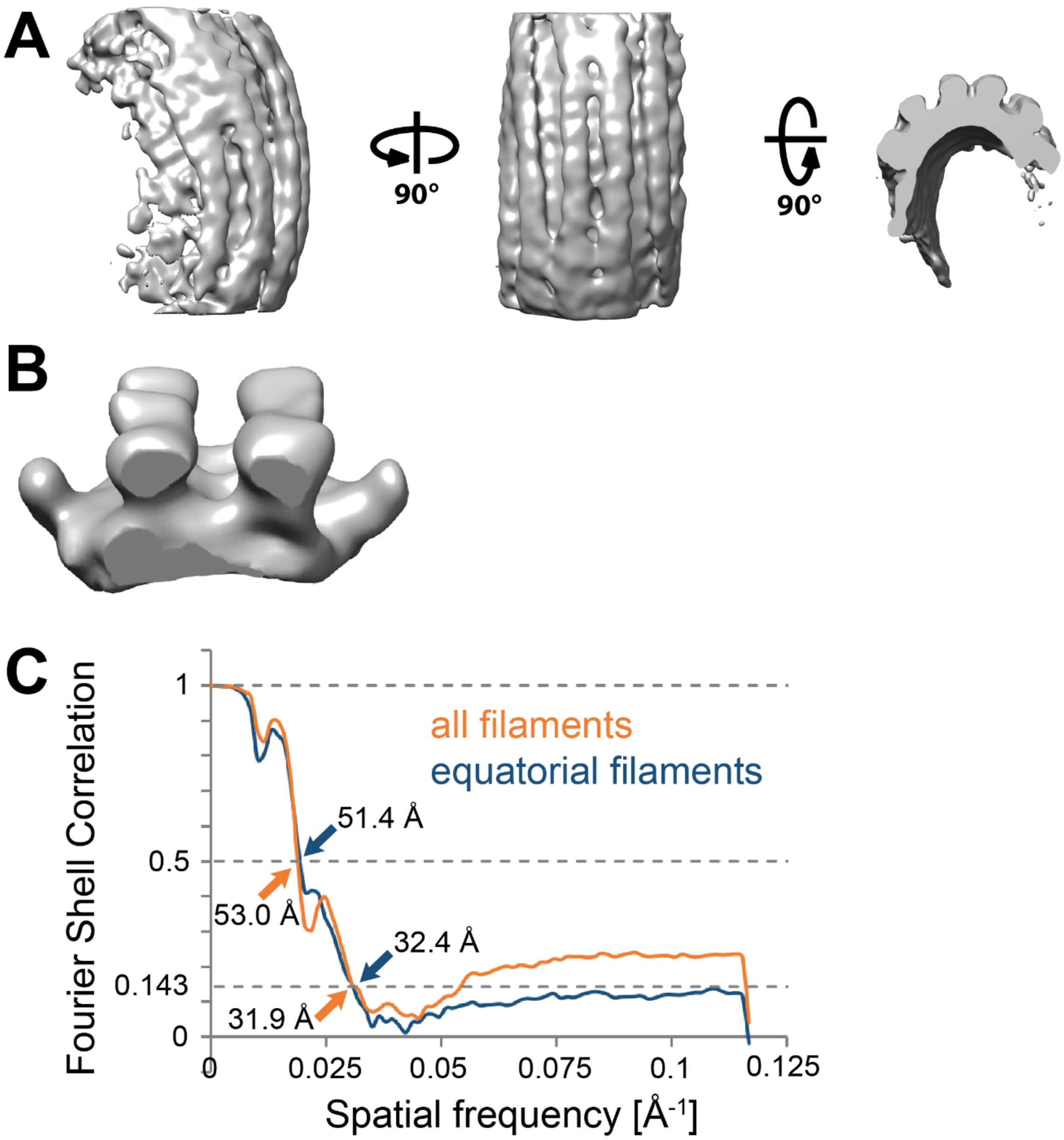
A
90°
90°
B
C
Fourier Shell Correlation
1
0.5
0.143
0
0
0.025
0.05
0.075
0.1
0.125
Spatial frequency [Å-1]
all filaments
equatorial filaments
51.4 Å
53.0 Å
32.4 Å
31.9 Å

**Fig. S2: ESCRT-III filament bundles form distinct clusters on the surface of helical tubes.** (A) Side view (left), top view (centre) and cross-section (right) of raw global subtomogram average showing filaments following the tube axis. (B) Side view of refined subtomogram average showing the detail of the equatorial filament cluster. (C) Fourier Shell Correlation for map shown in Fig. 2A-D and Fig. S2A. (D) Fourier Shell Correlations for maps shown in Fig. 2A-C and Fig. S2A (orange), and Fig. 2E-G and Fig. S2B (blue).

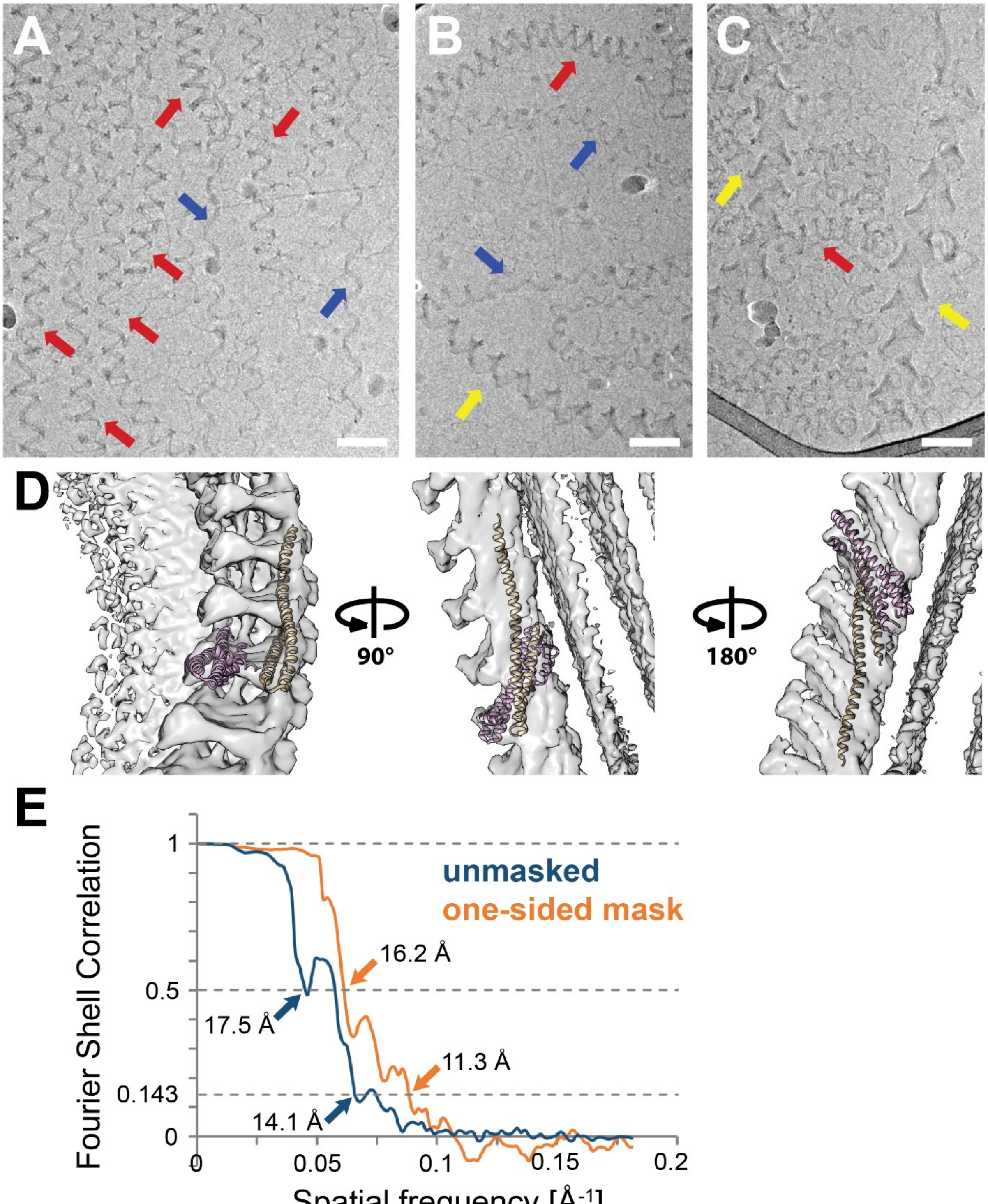
A
B
C
D
90°
180°
E
Fourier Shell Correlation
Spatial frequency [Å-1]
unmasked
one-sided mask
16.2 Å
17.5 Å
11.3 Å
14.1 Å
1
0.5
0.143
0
0
0.05
0.1
0.15
0.2

**Fig. S3: Organisation of tube-less ESCRT-III filaments.** (A-C) Electron micrographs showing different vitrified, tube-less, helical ESCRT-III filament bundles formed upon detergent removal. The major population of ribbons presents a zig-zag shape (red arrows; see also Fig. 3A, D), a second population appears sinusoidal (blue arrows; see also Fig. 3B, E) and a third, more complex helical ribbon filaments with higher substrand numbers (yellow arrows; see also Fig. 3C, F). Scale bars 100 nm. (D) Asymmetrically refined structure (see Fig. 2E) showing the result of docking an ESCRT-III subunit crystal structure in the open conformation (*D. melanogaster* CHMP4B homolog Shrub; PDB 5J45) (*11*), shown in orange, fitted in the outer strand of the filament. Fitted in the inner strand, a closed conformation ESCRT-III subunit is shown in pink (Human CHMP3; PDB 3FRT) (*12*). (E) Fourier Shell Correlations for maps shown in Fig. 3G (unmasked, blue) and Fig. 2H and Fig. S2D (masked, orange).

**Movie S1.**

Scanning through slices of tomogram and rotation of filtered and segmented tomographic volumes as shown in Fig. 1C-E.

**Movie S2.**

Scanning through slices of tomogram and rotation of filtered and segmented tomographic volumes as shown in Fig. S1 I-K.
